# Bulky glycocalyx drives cancer invasiveness by modulating substrate-specific adhesion

**DOI:** 10.1101/2023.08.03.551677

**Authors:** Amlan Barai, Niyati Piplani, V Gomathi, Mayank M Ghogale, Sushil Kumar, Madhura Kulkarni, Shamik Sen

**Author notes:** Université Paris Cité, CNRS, Institut Jacques Monod, Paris, France.

## Abstract

Majority of the eukaryotic cell surface is decorated with a layer of membrane attached polysaccharides and glycoproteins collectively referred to as the glycocalyx. While formation of a bulky glycocalyx has been associated with cancer progression, the mechanisms by which the glycocalyx regulates cancer invasiveness is incompletely understood. We address this question by first documenting sub-type specific expression of the major glycocalyx glycoprotein Mucin-1 (MUC1) in breast cancer patient samples and breast cancer cell lines. Strikingly, glycocalyx disruption led to inhibition of 2D motility, loss of 3D invasion and reduction of clonal scattering of breast cancer cells at the population level. Tracking of 2D cell motility and 3D invasiveness of MUC1-based sorted sub-populations revealed fastest motility and invasiveness in intermediate MUC1-expressing cells, with glycocalyx disruption abolishing these effects. While differential sensitivity in 2D motility is attributed to a non-monotonic dependence of focal adhesion size on MUC1 levels, higher MUC1 levels enhance 3D invasiveness via increased traction generation. In contrast to inducing cell rounding on collagen-coated substrates, high MUC1 level promotes cell adhesion and confers resistance to shear flow on substrates coated with the endothelial surface protein E-selectin. Collectively, our findings illustrate how MUC1 drives cancer invasiveness by differentially regulating cell-substrate adhesion in a substrate-dependent manner.

## Introduction

The cell interface is critical in terms of maintaining cell-substrate interactions. The cell senses its surroundings and responds accordingly using different receptors on the cell surface, of which perhaps the most important are a group of molecules that regulate cell-substrate adhesions. Integrins—which mediate cell-matrix adhesions—form cellular mechanosensory components that can regulate cytoskeletal architecture and control cellular dynamic properties including cell shape change and cell migration^1–4^. Mammalian cells are often coated with a layer of sugars mainly of glycoproteins and glycolipids in origin, which are collectively referred to as the glycocalyx^5^. The glycocalyx layer forms a coat around the cell; hence, most of the surface signal receptors including integrins and cadherins are buried in the glycan crowd. Therefore, change in the glycan layer is expected to significantly impact various surface signalling pathways. In addition, since glycocalyx is the outermost layer and is in direct physical contact with the cell surroundings, biophysical properties of the glycan layer should greatly impact cell-substrate interactions, and consequently, cell-substrate adhesions.

The glycocalyx layer shows a remarkable change in its composition and organization during development and disease like cancer^6–10^. Recent studies have demonstrated that malignant cancer cells often overexpress a group of transmembrane glycoproteins called mucins that make up most of the cell surface glycan structure commonly referred to as the bulky glycocalyx. The mucin family includes several trans-membrane proteins characterized by extensive glycosylation of PTS domains (proline, threonine and serine rich domain). Mucin 1 (hereafter referred to as MUC1) has been more extensively studied among other mucins and its overexpression has been documented in multiple cancers including breast cancer^10–13^. Instead of exhibiting an apical localization profile, MUC1 localizes all over the cell surface in cancer cells. These alterations in expression and localization may trigger loss of adhesions and increased invasiveness. Consistent with this, MUC1 transfection in human pancreatic and gastric cells led to decreased adhesion to type I collagen, type IV collagen, fibronectin, and laminin, but increased motility and in vitro invasiveness^14,15^. In contrast, inhibition of MUC1 in human pancreatic cancer cells led to slower proliferation, increased cell-matrix adhesion, and reduced lymph node metastasis^16–18^.

While these studies are indicative of an inverse correlation between MUC1 expression and cell-matrix adhesions, in a seminal study, Weaver and co-workers demonstrated that a bulky glycocalyx formed by overexpressing the extracellular domain of MUC1 can drive tumour progression through the formation of integrin-based adhesions and integrin-based signaling^19–21^. The bulkiness of MUC1 can create steric hindrance and induce integrin funnelling into pre-existing adhesions, thereby effectively increasing the size of adhesions^14–16^. Recent studies have also demonstrated significant contributions of the surface glycocalyx in cell membrane shape regulations and membrane protrusion formation^22,23^. Collectively, these studies suggest that MUC1 regulates invasiveness by modulating cell-matrix adhesion. However, how such alterations impact different stages of invasion (e.g., invasion through the stroma, or in circulation) remains unclear.

This study investigates glycocalyx alterations associated with cancer, and their biophysical role in regulating cell-substrate adhesions and cancer invasiveness. We found altered glycocalyx expression and localization differentially regulate the cell-substrate adhesion in a substrate-dependent manner. With increasing mucin levels, cell initially transforms from epithelial to mesenchymal state, and further increase help them to transform to a circulating tumour cell-like stage.

## Materials and methods

### Cell culture

MCF7, ZR-75-1, and MDA-MB-231 cell lines were obtained from the National Center for Cell Science (NCCS), Pune, India. MCF7 and MDA-MB-231 cells were grown in high glucose DMEM (Invitrogen, Ref # 11965084) and ZR-75-1 were cultured in RPMI-1640 (Gibco, Ref# 31800-022). Media was supplemented with 10% fetal bovine serum (FBS, HiMedia, Cat # RM9952). MCF 10A cells were obtained from ATCC (Cat# CRL10317) and were grown in mammary epithelial growth media (Cat# CC-3150, Lonza) supplemented with cholera toxin (Cat# C8052, Sigma). Cells were maintained at 37 °C and 5% CO_2_ levels and passaged at 70–80% confluency using 0.25% trypsin-EDTA (HiMedia, Cat # TCL099). For experiments, substrates were coated with 10 μg/cm^2^ collagen type-I from rat tail (Sigma, Cat # C3867) or 0.2ug/mL E-selectin (Cat# 724-ES, R&D Systems) overnight at 4°C.

### Enzyme and drug treatment

Cells were treated with Neuraminidase (Sigma, Cat# N2876) to remove surface glycocalyx and tunicamycin (Sigma, Cat# T7765) was used to prevent new glycosylation^24,25^. Cells were treated with different concentrations of Neuraminidase with 20 µg/ml Tunicamycin for up to 36 hours to check cell viability by MTT assay (Supp. Fig 2). Based on the assay, 500 mU/ml Neuraminidase containing 20 µg/ml tunicamycin was used for all the experiments. Cells were treated with the enzyme cocktail for two hours prior to all experiments. For motility assays, enzyme cocktail was kept in the media during live cell imaging for up to 14 hours.

### FACS sorting

Upon reaching 60-70% confluency, cells cultured in tissue culture flasks were trypsinized. The pellets were resuspended in FACS (1% FBS in PBS) buffer and split into two tubes for stained and unstained (control) samples. Cells were stained with FITC conjugated anti Human MUC1 antibody (BD, Cat # 559774) for 30 minutes at 4 °C in dark condition at recommended dilution. Cells were centrifuged after staining and pellets were resuspended in FACS buffer (wash) and centrifuged again. Pellets were dissolved again in FACS buffer and passed through cell strainer to remove clumps. Prepared samples were then analyzed or sorted in sterile FACS tubes for further experiments using a BD FACS Aria^TM^ III Cell Sorter.

#### Single cell biophysical measurements

##### 2D & 3D motility

Cells were sparsely seeded (2000 cells/cm^2^) on collagen-coated 48-well cell culture plates. 12-14 hrs after seeding, motility videos were captured for 12-14 hrs at 15 minutes frame interval in a live cell imaging chamber (OKO Lab) using an Olympus inverted microscope (Olympus IX83). Cell trajectories and speed were measured from obtained videos using the manual track plugin in Fiji-ImageJ.

For 3D-motility studies, 3D-Collagen gels were prepared from rat tail collagen type-1 (Corning, Ref# 354249) stock solution accordingly the manufacturers protocol by mixing required amount of collagen with 10x PBS and cell culture media, and then adjusting the pH to 7.3 using 1 N NaOH solution. Cells harvested from cell culture dishes were mixed with precursor collagen gel solution at a density of 18,000 cells/ml such that the final concentration of the solution was 1.25 mg/ml. 180 μl of this collagen-cell mixture was dispensed in each well of a glutaraldehyde functionalized 48-well cell culture plate and incubated in a 37 °C incubator. Once gel formation, complete media supplemented with appropriate enzyme/drug/vehicle was gently poured on top of the gels and incubated in CO_2_ incubator for another 12 hrs prior to imaging. 3D motility videos were captured and analyzed the same way as described above for the 2D motility experiment.

##### (3)#Spheroid invasion and clonogenic assay

Spheroids generated by hanging drop method^26^ were embedded in 3D collagen gels and imaged every 24 hrs using an inverted microscope. Spheroid spread area was calculated and compared to the initial spheroid size from the acquired images. For clonogenic assay, cells were seeded at a very low density (∼800 cells/cm^2^) and were grown for 6 – 8 days to enable micro-colony formation. Cell microcolonies were then imaged using an inverted microscope. Colony features like cell-cell distance were measured using ImageJ.

##### Adhesion & De-adhesion assays

Cells from different conditions (FACS sorted, enzyme treated etc.) were seeded on Collagen or E-selectin coated 96-well plates. Equal number of cells were seeded in wells and incubated for 30 minutes at 37° C. After gentle washing with warm PBS, cells were fixed with 4% PFA for 20 minutes and stained with DAPI for visualizing the nuclei. Entire wells were then imaged using an inverted microscope and the number of attached cells/well was counted from obtained images using ImageJ.

De-adhesion assay was performed using protocol described elsewhere^27,28^. Briefly, cells grown on collagen/E-selectin coated dishes for 24 hrs were washed with PBS and incubated with warm (37°C) trypsin. Movies were acquired till cells rounded up but remained attached to their underlying substrates. De-adhesion timescales for individual cells were calculated from the acquired movies.

##### Focal adhesion (FA) Dynamics

For probing focal adhesion dynamics, mCherry Paxillin transfected cells were seeded on collagen-coated glass bottom dishes and imaged at 30 sec intervals for 30-45 min at 63× magnification using a scanning probe confocal microscope (Zeiss, LSM 780)^29^. Focal adhesion lifetime was analyzed from acquired normalized image sequences using protocols as described elsewhere^30^.

### Immunostaining

For immunostaining, cells were cultured on ligand-coated coverslips for 16 – 22 h prior to fixing with 4% paraformaldehyde. Fixed cells were permeabilized in 0.1% Triton X-100 for 8 – 12 min, blocked using 5% BSA for 2 h at room temperature, and then incubated with primary antibodies (anti-MUC1 (CD227) Mouse monoclonal (BD, Cat# 555925), anti-paxillin rabbit monoclonal antibody (Abcam, Cat # ab32084), anti phospho-focal adhesion kinase (pFAK) rabbit monoclonal antibody (Cat# sc-374668, Santa Cruz Biotechnology), anti MMP2 (Cat # 436000, Invitrogen) diluted in PBS overnight at 4°C. Coverslips were then washed with PBS and incubated for 1.5 hrs at room temperature (RT) with appropriate secondary antibodies diluted in PBS. After labelling nuclei with Hoechst 33342 (Cat# B2261, Merck) for 5 min at RT, coverslips were mounted using mounting media. For lectin staining, FITC conjugated Lectin from Triticium vulgaris (FITC-WGA) (Sigma, Cat# L4895) was used. Mounted coverslips were imaged using a scanning probe confocal microscope (LSM 780, Zeiss) using 63× objective. Images were processed and analyzed using Fiji-ImageJ software.

### Electron microscopy

For scanning electron microscopy experiments, cells were grown on ligand coated glass coverslips or on top of 3D collagen and were then fixed with modified Karnovsky’s fixative (4% PFA and 2% glutaraldehyde in PBS) initially for 20 minutes at RT and then at 4°C for 24 – 48 hrs. After further fixation with 1% Osmium tetroxide, cells were dehydrated using 30%/50%/ 70%/90%/100% ethanol in a step-wise manner with each incubation lasting 5 – 10 min. After a final drying with hexamethyldisilazane (HMDS), samples were mounted on a stab, sputter coated using PP3000T (Quorum Technologies) and imaged at appropriate magnification using Joel JSM-7600F scanning electron microscope.

### Microfluidics experiments

For shear experiments, a straight channel 400 μm in width with 2 ports was designed (Fig. 7G). After patterning the design on a silicon wafer using photolithography with SU-8 photoresist (MicroChem), PDMS (Sylgard 184) was poured onto the master, and cured in the oven. After peeling, the PDMS substrates were plasma bonded to glass bottom petri-dishes and 2 ports were punched using biopsy punch. Devices were coated with 0.2 µg/ml E-selectin overnight at 4°C and blocked with 5% BSA solution for 1 hr at RT prior to seeding cells and allowing them to adhere for 1 hr. The device was connected to a syringe pump and placed under an inverted epifluorescence microscope with 37°C stage incubator. Cells were subjected to varying flow rates using the pump and live cell movies of the entire device were recorded at 4× using tile imaging. Live cell movies were also obtained using 60× oil immersion objective after staining the cells with FITC-WGA (30 min incubation with 1× FITC-WGA in the media).

RT PCR: Total RNA was eluted from knockdown and control cells using the RNeasy® Plus mini kit (Qiagen, Ref # 74134). 2 µg of total RNA was reverse transcribed using High Capacity cDNA Reverse Transcription Kit (AppliedBiosystems, Ref. # 4368814) and amplified using PowerUp^TM^ SYBR Green Master Mix (AppliedBiosystems, Ref. # A25742) in a QuantStudio 5 (AppliedBiosystems). PCR data were analyzed using comparative Ct method and Cyclophilin A expression was used to normalize gene expression data. Primers used for RT-PCR were Mucin-1-Forward: 5’-GCCTCTCCAATATTAAGTTCAG, Mucin-1-Reverse: 5’-AGTCGTCAGGTTATATCGAG, Cyclophilin A -Forward: 5’-TGGGCCGCGTCTCCTTTGA, and Cyclophilin A -Reverse: 5’-GGACTTGCCACCAGTGCCATTA.

Immunohistochemistry of patient samples and scoring: IHC staining was performed on formalin-fixed paraffin embedded (FFPE) tissue. 3 µm sections were obtained with a microtome (Leica RM2255 Fully Automated Rotary Microtome). The sections were transferred onto positively charged slides, dried at 62°C overnight, and deparaffinized and hydrated using xylene and ethanol dilutions. For IHC, the Epredia™ UltraVision™ Quanto Detection System HRP DAB (Fisher scientific, TL015QHD) was used. After incubation with Anti-MUC1(BD Pharmingen, 555925) at 4°C overnight, immunodetection was performed with 3,3’-diaminobenzidine chromogens available in the kit. Imaging of all slides was done by OptraScan – OS-15 bright field digital scanner at 400X magnification. Images were viewed using the ‘Image scope’ software and processed for scale bar. MUC1 expression was scored according to the percentage of tumor cells positive for MUC1 staining.

### Copy number alteration (CNA) analysis

Copy number alterations of MUC1 family members were analysed using two patient datasets (METABRIC and Firehose Legacy) by cBioPortal^31^. Individually, either METABRIC or Firehose Legacy was selected for analysis.

Analysis of single-cell RNA-sequencing data: For breast cancer CTCs, the count matrix and single-cell experiment object were obtained from the GEO accession number GSE109761^32^. The authors provided an object (GSE109761_sce_hs.rds.gz) containing CTCs from 34 patients and patient-derived xenografts (PDXs). We downloaded the object and transformed the Fragments Per Kilobase of transcript per Million mapped reads (FPKM) counts into Transcripts Per Million (TPM) counts. The expression levels of MUC1 in different clusters of CTCs were visualized using the Matplotlib library.

For the primary tumor cells, the count matrix and single-cell experiment object for primary breast cancer tumor were obtained from the GEO accession number GSE118390^33^. Epithelial cells were filtered from the count matrix using the clustering code provided by the authors. The cluster with a maximum number of epithelial cells was taken for analysis. The average expression of MUC1 in this cluster was then calculated using the function provided by the authors. The obtained expression was then plotted as a line plot using Matplotlib.

### Statistical analysis

Data distribution was tested using the Kolmogorov-Smirnov normality test. Based on the outcome, either parametric or nonparametric statistical test was performed. For parametric data, statistical analysis was performed using one-way ANOVA and Fisher’s test was used to compare the means. Mann-Whitney test was performed for non-parametric data. Statistical analysis was performed using Origin 9.1 (OriginLab® Corporation), with p <0.05 considered to be statistically significant.

## Results

### Mucin 1 (MUC1) expression and/or glycosylation are elevated in breast cancer

Aberrant glycosylation and upregulation of bulky glycocalyx have already been reported in various cancer cells including pancreatic cancer, lung cancer, ovarian cancer, bladder cancer and prostate cancer (CITE relevant papers against each cacner type). Copy number alterations (CNAs) encompass gene amplifications, gains, deep or shallow deletions, play a crucial role in cancer development and progression. We performed CNA analysis of mucin genes to identify genomic alterations present across the breast cancer. Oncoprint represented the distribution of CNAs in mucin (Supp. Fig. 1). Noticeably, the MUC1 gene had the highest probability in genomic amplifications (21% in METABRIC and 12% in Firehose Legacy) among mucins, suggesting a potential oncogenic role of MUC1. Our immunohistochemical staining revealed that ER+ and HER2+ breast tumors have high mucin expression; especially, the aggressive HER2+ breast cancer shows significantly more MUC1 positive fraction compared to triple negative breast cancer (Figs. 1A, B). However, MUC1 expression was not significantly increased in ER+ tumor when compared with ER-tumor (Fig. 1C).

**Figure 1:**
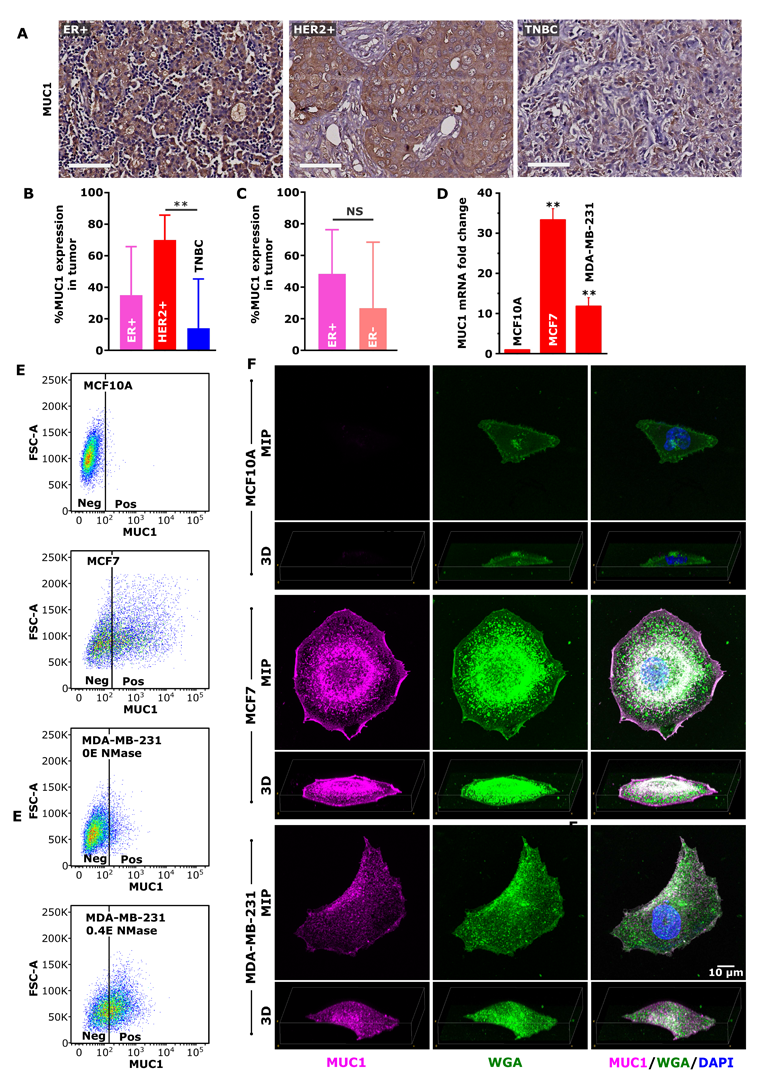
Cell surface bulky glycocalyx is upregulated in cancer. (A) Representative images of MUC1 immunohistochemical staining in ER+, HER2+ and triple negative breast cancer (TNBC) tissue. Scale bar 50 µm. Quantification of MUC1 expression from immunohistochemical staining from ER+, HER2+, and TNBC staining (n = 5 in each case) in (B) and ER+ (n = 6) and ER-(n = 9) samples in (C). ** p < 0.005, ns p > 0.05, data is presented as mean ± SD). (D) Analysis of MUC1 mRNA transcripts using real-time PCR. Total RNA harvested from 24 h culture of MCF10A, MCF7 and MDA-MB-231 cells were subjected to quantitative real-time PCR analysis (n≥3, **P<0.001; values indicate mean ± SEM). (E) FACS analysis of cell surface MUC1 expression in MCF10A, MCF7 and MDA-MB-231 cell population. (F) Confocal microscopy images showing surface glycan distribution in MCF7 and MDA-MB-231 cells. Unpermeablized cells were stained with MUC1 antibody and FITC-WGA (Lectin) and then glycocalyx was visualized from maximum intensity projection (MIP) and reconstituted 3D image. F-actin was stained with phalloidin and nucleus was stained with DAPI. Scale bar 10 µm.

Next, we assayed MUC1 expression in breast cancer cell lines. In comparison to non-malignant MCF10A cells, MUC1 expression was upregulated in both MCF7 and MDA-MB-231 (hereafter referred to as MDA231) cells (Fig. 1D). Consistent with transcript profiles, FACS analysis revealed mostly MUC1 negative (MUC1-neg) cells in MCF10A cells (Fig. 1E). Interestingly, considerable heterogeneity in MUC1 expression was observed in MCF7 cells with the presence of MUC1-Low and MUC1-High expressing cells, as well as a small subpopulation of MUC 1 negative cells. While FACS staining revealed the presence of MUC1 negative and MUC1-Low sub-populations in MDA-231 cells, substantial increase in the proportion of MUC1 positive cells upon neuraminidase (NMase) treatment suggests that increased MUC1 glycosylation in MDA-MB-231 cells is restrictive for antibody to stain MUC1, as reported elsewhere^34^. Confocal imaging further confirmed the presence of MUC1 on the cell surface of MCF7 and MDA231 cells and its absence in MCF 10A cells. To test overall glycocalyx levels in these three cell-types, we stained cells using wheat germ agglutinin (WGA), a lectin that largely stains sialic acid and associated sugars. The staining revealed that both MCF7 and MDA231 have an overall more glycosylated surface compared to MCF 10A (Fig. 1F). Together, these results reveal increased MUC1 levels and its glycosylation in breast cancer cells compared to non-transformed cells.

### Enzymatic de-glycosylation of cell surface glycocalyx reduces cancer invasiveness

The sugar residues on the glycoprotein provide stability and hence the removal of sugar leads the glycan layer to collapse. Neuraminidase (NMase) from Clostridium perfringens partially de-glycosylates the glycan layer thereby reducing the surface glycan layer thickness (Fig 2A, B). Enzyme treatment up to 36 hrs showed no significant impact on cell viability as checked via MTT assay, hence all the experiments were performed before 36 hr timepoint (Supp. Fig. 2).

**Figure 2:**
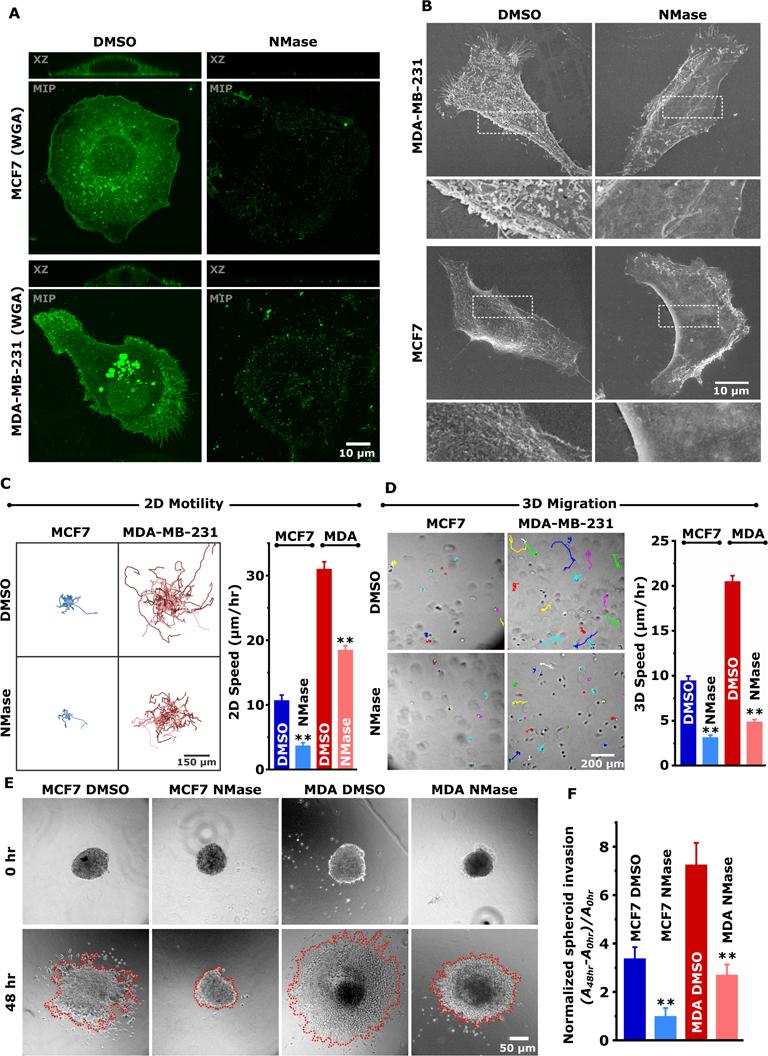
Enzymatic de-glycosylation of cancer cell surface glycocalyx reduces malignancy. (A) Confocal microscopy images showing maximum intensity projection (MIP) and XZ projection of FITC-wheat germ agglutinin (WGA) stained MCF7 and MDA-MB-231 cell after 3 hr DMSO or 0.4 U/ml Neuraminidase (NMase) treatment. (B) FEG-SEM images showing membrane microarchitecture of untreated (DMSO) and Neuraminidase treated (0.4 U/ml NMase) MCF7 and MDA-MB-31 cells. (C) (Left) Representative trajectories of MCF7 and MDA-MB-231 cells migrating on collagen-coated cell culture plate in the presence of 0.4 U/ml neuraminidase and 20 µg/ml Tunicamycin (NMase) or vehicle (DMSO). (Right) Quantification of 2D speed (n=3, per condition >70 cells, ** p < 0.005, data is presented as mean ± SEM). (D) (Right) representative microscopic frames showing cells encapsulated in 1.5 mg/ml 3D collagen gel with movement trajectories of MCF7 and MDA-MB-231 cells with NMase or DMSO treatment. (Left) Quantification of 3D speed (n=3, per condition >110 cells, ** p < 0.005, data is presented as mean ± SEM). (E) Representative phase-contrast images showing spheroid invasion assay in 3D collagen. Spheroids prepared from MCF7 and MDA-MB-231 cells were embedded in 3D-collagen layer in presence NMase or DMSO treatment and images were taken at 0 hr and 48 hr time points. (F) Quantification of spheroid invasion (n=3, 5-10 spheroids per condition, ** p < 0.005, data is presented as mean ± SEM).

To test the role of glycocalyx on cell invasiveness, we checked cell migration using live-cell migration setup in MCF7 and MDA231 cells (Fig. 2C). Deglycosylation with NMase treatment led to a significant reduction in 2D cell motility in both MCF7 and MDA-231 cells. To test whether a similar phenomenon is observed in the 3D scenario, we encapsulated cells in collagen hydrogels that more closely mimic the *in vivo* stroma and checked cell invasiveness after glycan disruption. In 3D also, the cells showed a similar response where the cell invasiveness dropped significantly upon neuraminidase treatment (Fig. 2D). In comparison, non-malignant MCF 10A cells did not show any significant change upon neuraminidase treatment (Supp Fig. 3). In line with these observations, enzymatic de-glycosylation also led to decreased spheroid invasion in MCF7 and MDA231 cells (Figs. 2E, F). Together, these observations suggest that glycan removal negatively impacts migration and invasion of cells while noncancerous cells remain unimpacted.

**Figure 3:**
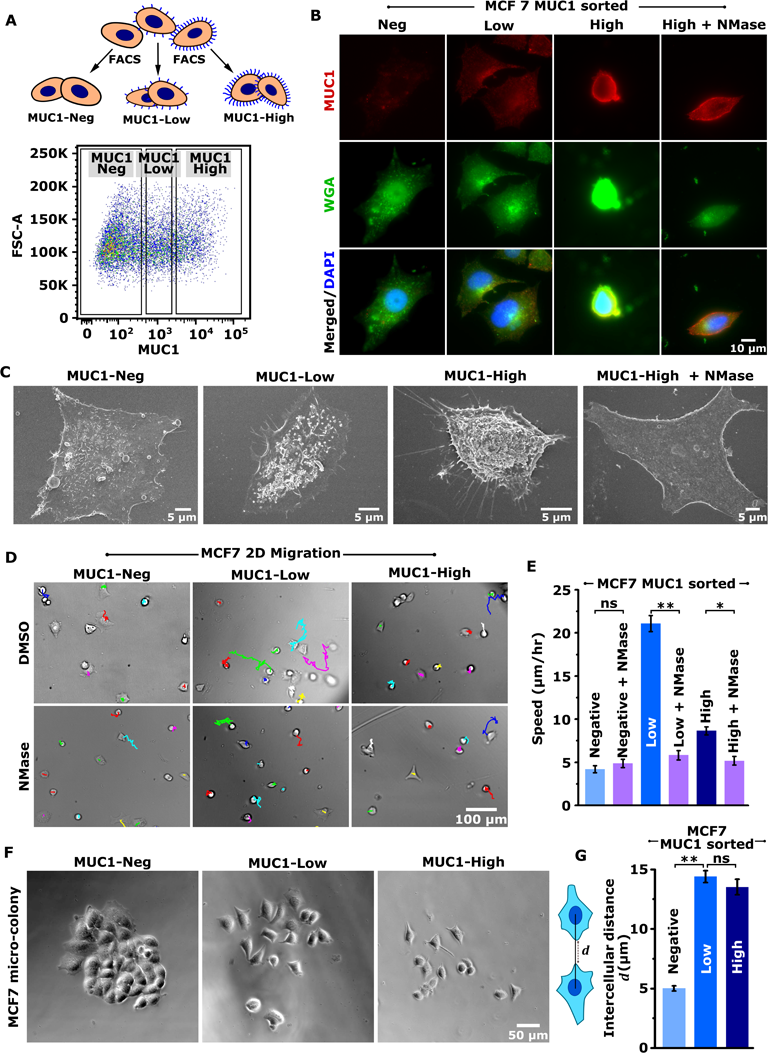
Intermediate level of glycocalyx promotes malignancy. (A) FACS sorting of MCF7 in mucin-1 negative, intermediate (low) and high cell populations. (B) Mucin 1 (MUC1) and FITC-wheat germ agglutinin (WGA) staining of FACS sorted MC7 cells. High mucin 1 expressing cells was also de-glycosylated with neuraminidase treatment (High + NMase). Scale bar 10 µm. (C) FEG-SEM images showing membrane microarchitecture of FACS-sorted MCF7 cells. Scale bars 5 µm. (D) 2D motility of FACS sorted MCF7 cells on collagen-coated substrate with images showing representative tracts and (E) quantification of cell speed. Scale bar 100 µm (n=3, 100-150 cells per condition, ns = non-significant, * p < 0.05, ** p < 0.005, data is presented as mean ± SEM). (F) Clonogenic assay showing micro-colony architecture of FACS sorted MCF7 cells. FACS-sorted cells were sparsely seeded and grown for 96 hrs so that cells can divide to form micro-colonies. The bottom graph measures the average intercellular distance between neighbouring cells in each colony. Scale bar 50 µm. (n=3, 30-40 colonies per condition, ns p > 0.05, ** p < 0.005, data is presented as mean ± SEM).

### Intermediate level of glycocalyx promotes invasiveness by regulating cell-matrix adhesions

Thus far, we have established that the presence of a glycocalyx increases cancer cell migration. To test whether different levels of surface glycocalyx differentially regulates cell migration, we have FACS-sorted MCF7, which exhibited a wide range of glycan distribution, into three sub-populations based on MUC1 expression (Fig 3A). Lectin staining of sorted MUC1 negative cells (MUC1-Neg), low to intermediate MUC1 expressing cells (MUC1-Low), and high MUC1 expressing cells (MUC1-High) revealed a close correlation between overall glycocalyx expression and MUC1 expression (Fig 3B). In comparison to the flattened morphologies of MUC1-Neg cells, MUC1-Low and MUC1-High were comparably rounded, with NMase treatment of MUC1-High cells leading to flattening similar to that of MUC1-Neg cells (Fig 3C).

Motility experiments on collagen coated dishes revealed that MUC1 positive (i.e. MUC1-Low and MUC1-High) cells migrate faster than MUC1-Neg cells. Surprisingly, MUC1-Low cells were found to migrate several times faster than MUC1-High cells (Figs. 3D, E). This differential migratory phenotype can be attributed to the surface glycocalyx expression as enzymatic deglycosylation reduced migration speed to that of MUC1-Neg cells across all conditions (Figs. 3D, E).

When cells were seeded sparsely and allowed to form microcolonies, MUC1-Neg cells were found to form compact epithelial-like colonies with low intercellular distance, while MUC1 positive cells formed sparse colonies with greater intercellular distance resembling a more mesenchymal phenotype (Fig 3F, G). To check if the cell morphology is impacted by surface glycan level, we have analyzed single cell morphology of sorted cells grown on collagen coated surface from phase contrast images. While cell spread area reduced with increase in MUC1 levels, enzymatic deglycosylation led to increase in cell spread area to levels comparable to that of MUC1-Neg cells (Figs. 4A, B). MUC1-High cells also possessed the highest circularity which dropped to baseline levels after neuraminidase treatment (Figs. 4A, C).

**Figure 4:**
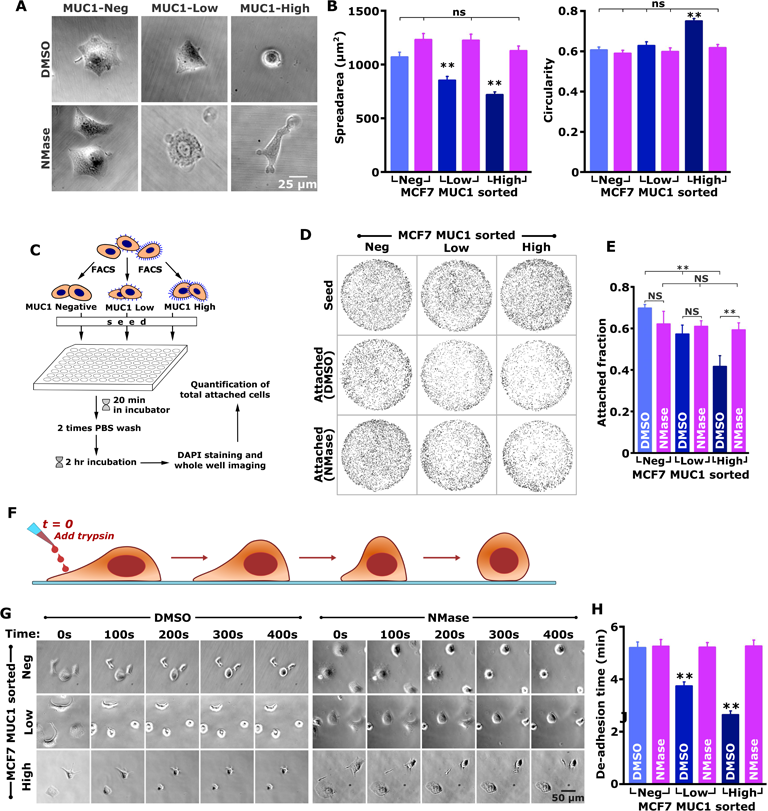
Intermediate level of glycocalyx promotes malignancy by regulating focal adhesion. (A) Representative phase contrast images showing morphology of MUC1 FACS sorted MCF7 cells cultured on collagen-coated substrate. MCF7 cells are sorted in mucin-1 negative, intermediate (low) and high MUC1 expressing cell populations and were grown for 14 hours in presence of DMSO control or 0.4 U/ml Neuraminidase and 20 µg/ml tunicamycin (NMase). (B) Quantification of cell spread area and cell circularity form obtained images (n>100 cells from 3 independent experiments, ns p > 0.05, ** p < 0.005, data is presented as mean ± SEM). (C) Schematic of adhesion assay with MUC1 FACS sorted MCF7 cells. Facs sorted MCF7 cells are seeded in collagen coated 96-well plate and were allowed to attach for 20 minutes, was then PBS washed and was stained with DAPI and the whole plate was imaged using microscope to count cells. (D) Representative images of nucleus in one well after thresholding with top row showing wells without wash step (seed) and the bottom panel is showing attached cells with and without NMase treatment. (E) Quantification of attached 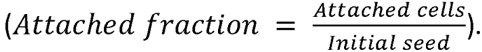 (n=3, ** p < 0.005, ns p > 0.05, data is presented as mean ± SEM). (F-H) De-adhesion assay of MUC1 FACS sorted MCF7 cells without and with enzymatic deglycosylation using neuraminidase treatment (NMase). (F) Schematics of the de-adhesion assay. (G) Representative frames showing cell de-adherence over time and (H) Graph is showing average de-adhesion time (n=3, 90 - 150 cells per condition, ** p < 0.005, NS p > 0.05, data is presented as mean ± SEM).

Alterations in cell spread area hints towards a differential regulation of cell-substrate adhesion by the cell surface glycocalyx. To test whether cell-substrate adhesion is altered by MUC1 levels, we performed cell adhesion assay wherein MUC1 sorted cells were allowed to adhere for 2 hrs on collagen-coated dishes, and then subjected to PBS wash (Fig. 4C). Nuclei-based counting revealed an inverse correlation between MUC1 levels and the proportion of attached cells (Figs. 4C-E). To further test how MUC1 levels influence adhesion, trypsin de-adhesion assay^28,35^ was performed wherein spread cells were incubated with warm trypsin and their detachment kinetics tracked for the duration cells rounded up but remained attached to their substrates (Figs 4F, G). Once again, the de-adhesion time, i.e., the time for cells to round up scaled inversely with MUC1 levels with fastest de-adhesion observed in MUC1-High cells, and neuraminidase treatment abolishing the MUC1-dependent behavior (Fig. 4H).

Adhesion and de-adhesion experiments together suggest that glycocalyx regulates cell-substrate adhesion. To further test the molecular mechanism, we stained focal adhesions of sorted cells using paxillin staining (Fig. 5A). In comparison to small-sized focal adhesions observed in MUC1-Neg cells observed both at the cell periphery and the cell interior, prominent peripheral focal adhesions were observed in MUC1-Low cells (Figs 5A, B). Strikingly, in MUC1-High cells, few small-sized focal adhesions were detected. This MUC1-dependent focal adhesion formation was abolished upon NMase treatment.

**Figure 5:**
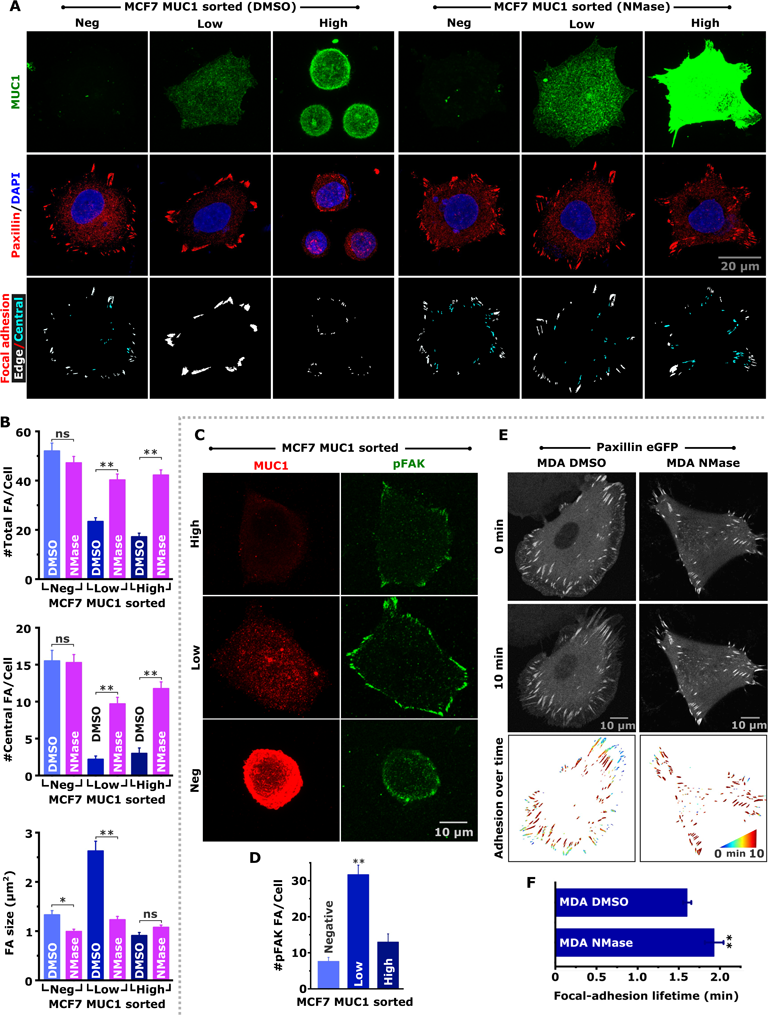
Glycocalyx regulates focal adhesion dynamics. (A) Representative confocal images of paxillin (red) and MUC1 (green) stained FACS sorted mucin-1 negative, intermediate (low) and high MCF7 cells in presence of DMSO control or 0.4 U/ml Neuraminidase and 20 µg/ml tunicamycin (NMase). Nuclei stained with DAPI. Bottom panel shows focal adhesion distribution with focal adhesion at the edge in white and adhesion inside in cyan. (B) Quantification of average number of total focal adhesion/cell, central focal adhesion/cell and average size of focal adhesions across different conditions (n=3, more than 50 cells per condition, ** p < 0.005, ns p > 0.05, data is presented as mean ± SEM). (C) Representative confocal images of phospho focal adhesion kinase (pFAK in green) and MUC1 (red) stained FACS sorted MCF7 cells and (D) Quantification of average number of pFAK focal adhesion/cell across different conditions (n=3, per condition ≥ 40 cells per condition, ** p < 0.005, ns p > 0.05, data is presented as mean ± SEM). (E) Focal adhesion dynamics in MDA-MB-231 cells in presence of DMSO control or 0.4 U/ml Neuraminidase and 20 µg/ml tunicamycin (NMase). Representative images showing focal adhesion at the beginning (0 min) and end (10 min) of experiment, and color-coded images depicting images of adhesions overlaid from multiple time points acquired over a period of 10 min. (F) Analysis of focal adhesion lifetime (*P<0.05; **P<0.001; NS p > 0.05, for n ≥ 16 cells analyzed per condition from three independent experiments, data is presented as mean ± SEM).

Phospho-focal adhesion kinase (pFAK) staining revealed highest active focal adhesions in MUC1-Low cells compared to MUC1-Neg and MUC1-High cells, with pFAK clusters located peripherally (Figs. 5C, D). Active focal adhesion formation in MUC-Low cells was associated with increased MMP2 localization at the cell membrane, while the total MMP2 expression remained unaltered (Supp Fig. 4). Focal-addition dynamics of mCherry-paxillin transfected malignant MDA231 cells further revealed that enzymatic deglycosylation significantly reduces focal-adhesion turnover rate (Figs. 5E, F).

Together these experiments suggest that the surface glycocalyx regulates cell-matrix adhesions. An intermediate level of glycocalyx expression promotes integrin-based focal-adhesion that are large in size and are peripherally located—a hallmark of mesenchymal cells^36–40^.

### Bulky glycocalyx increase 3D invasion by increasing cell surface traction

To test how cell surface glycocalyx impacts cell migration in 3D scenario where the cell is surrounded by extracellular matrix, we have checked migration rate of MUC1 sorted cells embedded in collagen hydrogel. Interestingly both MUC1-Low and MUC1-High cells migrated significantly faster compared to MUC1-Neg cells, while enzymatic deglycosylation led to near complete arrest in the cell motility (Figs. 6A, B). Cell migration often depends on how effectively cells can couple their internal actin retrograde flow with the substrate^41^, which is often achieved by focal adhesions that generate traction and propel the cell forward. Cell substrate traction can also increase by other means such as substrate typography^41^. To test the hypothesis that surface glycocalyx can induce higher cell-substate traction through increased surface roughness, we performed scanning electron microscopy (SEM) of sorted cells and enzyme treated cells grown on collagen coated glass coverslips (Figs. 2B, 3C, 6C). SEM images revealed that cell surface roughness increases with increasing MUC1 level and enzymatic deglycosylation makes the cell surface smooth (Figs. 2B, 3C). Further, MCF7 MUC1-High cells produce numerous membrane protrusions that effectively increase cell surface roughness (Figs 6C). To test how these protrusions would impact cells in 3D-ECM, we seeded cells on top of collagen hydrogel and prepared sample for SEM imaging. SEM images revealed that the cell substrate interface of high mucin expressing cells are rougher with numerous protrusions entangled in the collagen network (Fig. 6D). Such entanglements with the ECM may serve as transient adhesions that propel the cell forward during 3D migration.

**Figure 6:**
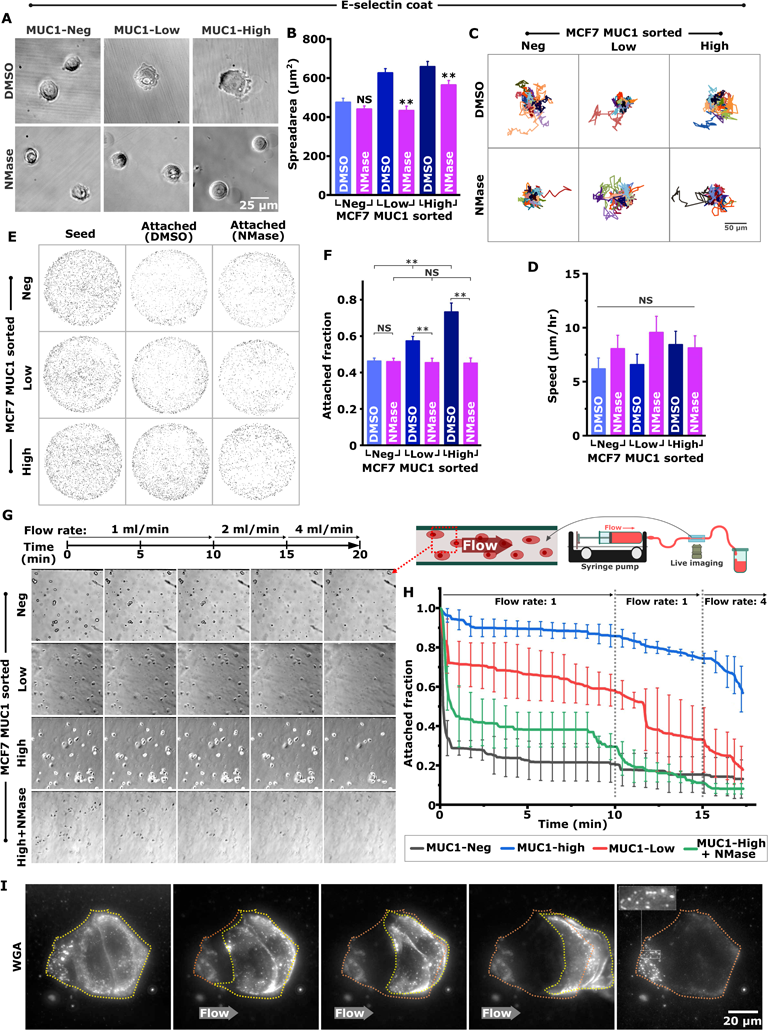
Bulky glycocalyx increase 3D invasion by increasing cell surface traction. (A-B) Invasion rate of FACS sorted MCF7 with mucin-1 negative, intermediate (low) and high MUC1 cells in 3D collagen gel (1 mg/ml) with (A) showing representative microscopic frames with migration tracts and (B) quantification of cell speed. Scale bar 100 μm. (n=3, 65-100 cells per condition, ** p < 0.005, data is presented as mean ± SEM). (C) FEG-SEM images showing membrane microarchitecture of FACS sorted MUC1-High MCF7 cells. Cells having high glycocalyx expression often shows numerous protrusions (white arrows) and blebs (blue arrow). (D) FEG-SEM images of FACS sorted MCF7 MUC1 negative and MUC1-High cells grown on collagen hydrogels. Right panel shows magnified inset with cell pseudo-colored in red shows more membrane micro-ridge (yellow arrow) and protrusions (green arrow) tangled in collagen hydrogels that likely increases membrane traction.

**Figure 7:**
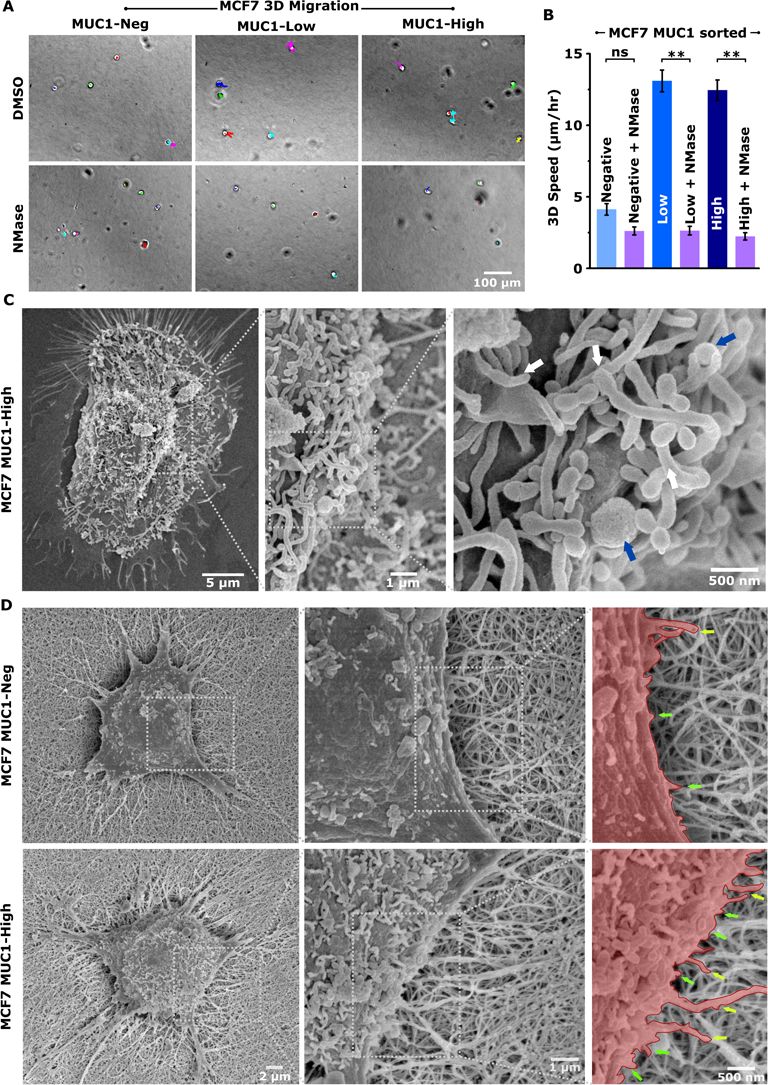
Bulky glycocalyx confers shear resistance and increased E-selectin adhesion. (A) Representative phase contrast images showing morphology of MUC1 FACS sorted MCF7 cells cultured on E-selectin coated substrate. MCF7 cells are sorted in mucin-1 negative, intermediate (low) and high MUC1 expressing cell populations and were grown for 14 hours in presence of DMSO control or 0.4 U/ml Neuraminidase and 20 µg/ml tunicamycin (NMase). (B) Quantification of cell spread area and cell circularity form obtained images (n>120 cells from 3 independent experiments, ns p > 0.05, ** p < 0.005, data is presented as mean ± SEM). (C) Representative trajectories of FACS sorted MCF7 cells migrating on E-selectin coated cell culture plate in the presence of 0.4 U/ml neuraminidase and 20 µg/ml Tunicamycin (NMase) or vehicle (DMSO). (D) Quantification of 2D speed (n=3, 55-70 cells per condition, ** p < 0.005, data is presented as mean ± SEM). (E) Adhesion assay with MUC1 FACS sorted MCF7 cells on E-selectin coated surface. Cells seeded on coated 96-well plate were allowed to attach for 20 minutes, was then PBS washed and was stained with DAPI and the whole plate was imaged using microscope to count cells. Representative images of nucleus in one well after thresholding with top row showing wells without wash step (seed) and the bottom panel showing attached cells with and without NMase treatment. (F) Quantification of attached cell 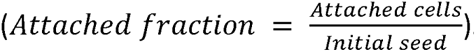 (n=3, ** p < 0.005, ns p > 0.05, data is presented as mean ± SEM). (G) FACS sorted MCF7 cells seeded on a selectin coated microfluidics channel were subjected to different flow induced shear stress and were imaged using a live cell imaging setup. Representative phase contrast images showing a section of the channel at different time points with attached cells (H) Quantification of remaining attached cell fraction across different conditions as they are subjected to flow induced shear stress (n≥3 data is presented as mean ± SEM). (I) WGA staining shows cell leave cell leaves glycan patch after detachment. WGA stained MCF7 cells seeded on a selectin coated microfluidics channel subjected flow induced shear was imaged using fluorescent live cell imaging setup. Yellow dotted line shows the cell boundary and orange dotted line shows initial cell adhesion area.

### Bulky glycocalyx confers shear resistance and increased E-selectin adhesion

Low adhesion and circular morphology of MUC1-High cells show striking similarities with circulating tumor cells (CTC) that are often found in the vasculature. To test if high levels of MUC1 can facilitate vascular metastasis, we have done experiments with E-selectin coated substrates as E-selectin coating mimics the vascular endothelial lining^42–44^. While cell spread-area decreases with increasing MUC1 level on collagen coated substrate (Figs. 4A, B), cells exhibited opposite response on E-selectin coated surfaces, i.e., with increasing MUC1 levels, cell adhesion increased on E-selectin coated substrates (Figs. 7A, B). This suggests that the glycocalyx promotes adhesion in a substrate-dependent manner. To test this further, we performed adhesion experiments on E-selectin coated plates with MUC1 sorted cells. This experiment also revealed increased attachment with increasing MUC1 levels, and neuraminidase treatment abolishing this dependence (Figs. 7E, F). Surprisingly, the altered adhesion did not significantly impact cell motility on E-selectin coated surfaces (Figs. 7C, D).

E-selectin coating alone does not fully recapitulate the vascular system in the absence of vascular fluid flow. To simulate the vascular system, we seeded cells on E-selectin coated channel (microfluidic device) connected to a syringe pump that can generate fluid (media) flow inside the channel. Live imaging at increasing flow rates revealed that with increase in MUC1 expression, cells take longer to detach while MUC1-Neg exhibited fastest detachment (Figs. 7G, H, Supp. Movie 1). Detachment of MUC1-High cells—which exhibited the strongest resistance to shear stresses— was significantly hastened upon enzymatic de-glycosylation. WGA-stained live imaging at high magnification revealed that cells shed glycocalyx and leave a trail of glycocalyx attached to the substrate (Figs. 6I, Supp. Movie 2). Cells are also known to regenerate surface glycocalyx^45,46^ (Supp. Figs. 5), thus suggesting a mode of rolling motility in the vasculature where E-selectin-glycan adhesions are broken by glycocalyx shedding from trailing edge and new adhesions are generated at leading edge from regenerated glycocalyx^47–49^ (Fig 8 schematic).

**Figure 8:**
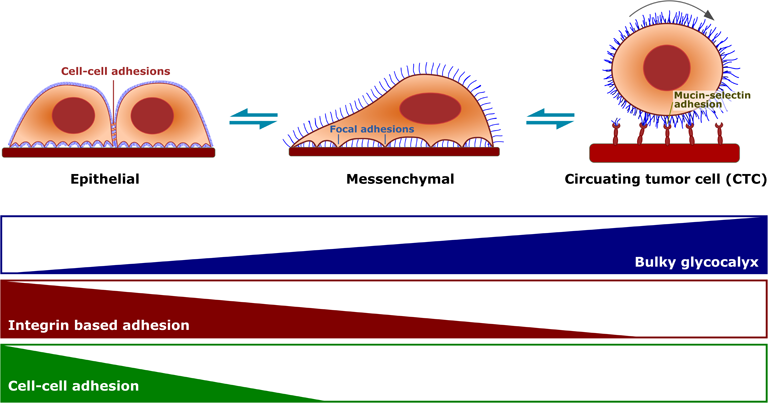
Schematic showing bulky glycocalyx mediated cancer progression. Cells can form diffused cell-substrate adhesion and retain intact cell–cell adhesion at low bulky glycocalyx level resembling epithelial cells. As bulky glycocalyx increases to an intermediate level, these cells transit to a mesenchymal state, where the cell loses cell-cell adhesion and gains clustered focal adhesion largely driven by glycocalyx-induced repulsion. Further glycocalyx upregulation causes a complete loss of integrin-based adhesion with high level of surface glycocalyx increases selectin based adhesion inside blood vessel and help in intravascular metastasis as circulating tumor cells.

Based on our *in vitro* data, we hypothesized that the expression of MUC1 in CTCs would be more heterogeneous compared to primary tumor cells. To understand the MUC1 expression in patient samples, we analysed single-cell RNA-sequencing data of patient CTCs and primary tumors of breast cancer. In support of our hypothesis, we observed that CTCs exhibit greater heterogeneity in MUC1 expression than primary tumor cells, as quantified by measuring the coefficient of variation, CV = 1.55 for CTCs and 1.09 for Primary tumors (Supp. Fig. 6). This enhanced heterogeneity in MUC1 expression in CTCs may contribute to their survival and resistance to the mechanical stresses encountered in the bloodstream.

## Discussion

Glycocalyx composition and organization changes significantly with cancer progression. Overexpression of several glycocalyx associated proteins including MUC1 has been reported in various cancer including breast cancer, lung cancer, pancreatic cancer, ovarian cancer, prostate and bladder cancer^6,13,50–53^. While MUC1 was long known to be associated with malignancy^12,13^, most of the mucin research was focused on its cytosolic domain which participates in various cancer-associated signalling cascade. The presence of bulky glycocalyx at the cancer cell surface is a rather recent finding^7^.

Here in this study, we demonstrate that bulky glycocalyx, on one hand being an effective spacer between cell membrane and the substrate, can physically regulate cell-substrate adhesion dynamics and integrin signalling. On the other hand, it can act as cell adhesion molecule that selectively binds to selectins that are abundantly present in the endothelial cells lining blood vessels and thus helps in vascular migration^42,54,55^. Additionally, glycocalyx can contribute significantly to cell membrane shape regulations and help membrane protrusion formation. The increased membrane micro-protrusions can increase membrane roughness and increase the cell-substrate traction force that increases cell migration. Since glycocalyx facilitates membrane protrusion and eases membrane bending, this would likely ease the production of protrusive structure like filopodia and lamellipodia formation.

Our initial experiments revealed that both MCF7 and MDA-MB-231 breast cancer cell surfaces are decorated with MUC1, and are overall more glycocalyx enriched compared to non-malignant MCF 10A cells. This higher expression of bulky glycocalyx increases cell migration and invasion that decreases upon enzymatic deglycosylation (Fig 2).

Cells exhibit diffused adhesions in the absence of a glycocalyx as bulky glycocalyx prevents adhesion formation by expanding the gap between the cell membrane and extracellular matrix (ECM)^19–21^. Cells in the presence of excess bulky glycocalyx, therefore, form integrin clusters at glycan excluded zones that forms clustered focal adhesions. Based on this, we hypothesized that optimal surface glycan levels enhance cell migration rate by optimally regulating cell-ECM adhesion, with the absence of glycocalyx hindering cell migration through increased adhesion formation, and excess glycocalyx preventing integrin-based adhesions altogether. Our experiment with FACS sorted MCF7 cells indeed confirmed that cells expressing intermediate MUC1 levels migrate faster than mucin negative and high MUC1 expressing cells (Fig 3D). Further investigation revealed this high cell migration speed is indeed caused by an optimal adhesion-deadhesion rate. Cells expressing an intermediate level of mucin have fewer focal adhesion and optimal focal adhesion turn-over compare to mucin negative cells which contribute to a faster migration. High mucin expressing cells are unable to form a sufficient adhesion to sustain adhesion-dependent migration (Figs 4, 5).

Circulating tumour cells are known to further up-regulate surface glycocalyx. It has been shown to have the highest level of glycocalyx expression. The cell surface glycocalyx can act as receptor molecules that can bind to selectins present on the surface of the vascular endothelial cells. Thus, the glycocalyx acts as adhesion molecules when the cancer cells are migrating through the blood vessels as circulating tumour cells. This is evident from the fact that high MUC1 expressing cells attach more strongly and spread more on selectin coated substrates compared to low MUC1 expressing cells (Figs 7 A-B, E-F). Additionally, on selectin coated microfluidic channels, high MUC1 expressing MCF7 cells were found to resist vascular fluid flow to the greatest extent and stay attached to the substrate (Fig 7G). Further, the cells might follow a rolling mode of migration that are often seen with leucocyte cells migrating through blood vessels^56,57^.

Therefore, based on our findings, we propose a mode of cancer metastasis dominated by surface glycocalyx expression: cells exhibit diffused adhesion with intact cell-cell adhesion at low bulky glycocalyx level resembling epithelial cells. With increasing glycocalyx, at intermediate levels, these cells transit to a mesenchymal state, where the cell loses cell-cell adhesion and gains clustered focal adhesions. Further upregulation causes the cell to lose their integrin-based adhesion altogether and prepares them for intravascular metastasis as circulating tumor cells that are largely mediated by selectin-glycan adhesion (Fig 8).

Apart from regulation of cell-matrix adhesions, glycocalyx has also been seen to regulate cell membrane microarchitecture^22,58^. Our SEM imaging experiments revealed the presence of micro-protrusions and small membrane blebs on the cell surface which disappears after enzymatic de-glycosylation. Further lack of these structures in MUC1 negative cells suggests that a bulky glycocalyx drives the formation of these membrane structures. Glycocalyx helps the formation of these micro-protrusions by decreasing membrane bending energy^22,23^. Glycan contributed membrane micro-protrusions lead to increased membrane roughness and generate membrane tractions that enable faster cell migration, as observed in our experiments (Figs 3C, 6). We believe these structures would additionally stabilize larger membrane protrusion in the 3D ECM network by tangling with the fibrous network and would form integrin independent adhesion (Fig 6D).

Overall, our study provided some interesting insights into the biophysical role of cell surface glycocalyx in cancer cell invasion, adhesion regulation and regulation of cell membrane biophysics.

## Supporting information

Supp. Figure 1 to 6, Supp.Movie1, 2 description

## Acknowledgments

SS acknowledges funding support from Department of Science and Technology, Ministry of Science and Technology (DST/SJF/LSA-01/2016-17), and IIT Bombay for providing Flow Cytometry (FACS) facility, Cryo FEG-SEM and Confocal Microscopy facilities. MK acknowledges the DBT-Ramalingaswami ‘re-entry’ fellowship and the DBT-Basic Research in Biology awarded by DBT-India and a research grant to CTCR by Bajaj Auto Ltd.

## Author Contributions

AB and SS conceived the study. AB, NP performed the cell experiments. GV and MK helped with the IHC. MMG and SK helped with the in-silico analysis. AB and SS wrote the manuscript.

---

**Supp. Figure 1:** Copy number alterations in mucin family genes in breast cancer tumor samples. OncoPrint visual of genomic alterations on query of mucin family members in METABRIC, n=2509 patients (A) and Firehose Legacy, n=1108 patients (B) using cBioPortal.

**Supp. Figure 2:** A dose-dependent response with varying concentrations of NMase and Tunicamycin showing treatment does not significantly affect cell viability. Cells were treated for 48 hrs and MTT assay was done to check viability (n=2, data is presented as mean ± SD).

**Supp. Figure 3:** (A) Representative microscopic frames showing cells entrapped in 1mg/ml 3D collagen gel with movement trajectories of MCF 10A, MCF7 and MDA-MB-231 cell in presence of 0.4 U/ml neuraminidase and 20 μg/ml Tunicamycin (NMase) or vehicle (DMSO). (B) Quantification of 3D speed (n=3, per condition >110 cells, ** p < 0.005, data is presented as mean ± SEM).

**Supp. Figure 4:** (A) Representative images showing total (maximum intensity projection, MIP) and bottom layer MMP2 levels in MUC1 FACS sorted MCF7 cells cultured on collagen coated substrate. MCF7 cells are sorted in mucin-1 negative, intermediate (low) and high MUC1 expressing cell populations and were grown for 14 hours prior to immunostaining. (B) Quantification of MMP2 expression from obtained image as bottom layer localization (top) and total level from MIP projection(bottom) (n = 3, ** p < 0.005, data is presented as mean ± SEM).

**Supp. Figure 5: Glycan recovery after enzyme treatment.** Cells were cultured for different duration after glycan removal with 0.4 U/ml Neuraminidase treatment to test their surface glycan recovery rate. (A) Representative WGA staining images showing surface glycan level at different time point after enzyme treatment. (B) Quantification surface glycocalyx recovery rate based on WGA intensity from obtained images (n = 2, ** p < 0.005, data is presented as mean ± SEM).

**Supp. Figure 6:** The line plots show the expression profile of the MUC1 gene in patient CTCs (A) and primary tumor cells (B).

**Supp. Movie 1:** FACS sorted MCF7 cells seeded on a selectin coated microfluidics channel were subjected to different flow induced shear stress and were imaged using a phase contrast live cell imaging setup.

**Supp. Movie 2:** WGA staining shows cell leave cell leaves glycan patch after detachment. WGA stained MCF7 cells seeded on a selectin coated microfluidics channel subjected flow induced shear was imaged using fluorescent live cell imaging setup using 60x oil immersion objective.

