## Supplementary material for "Bulky glycocalyx drives cancer invasiveness by modulating substrate-specific adhesion": Supp. Figure 1 to 6, Supp.Movie1, 2 description

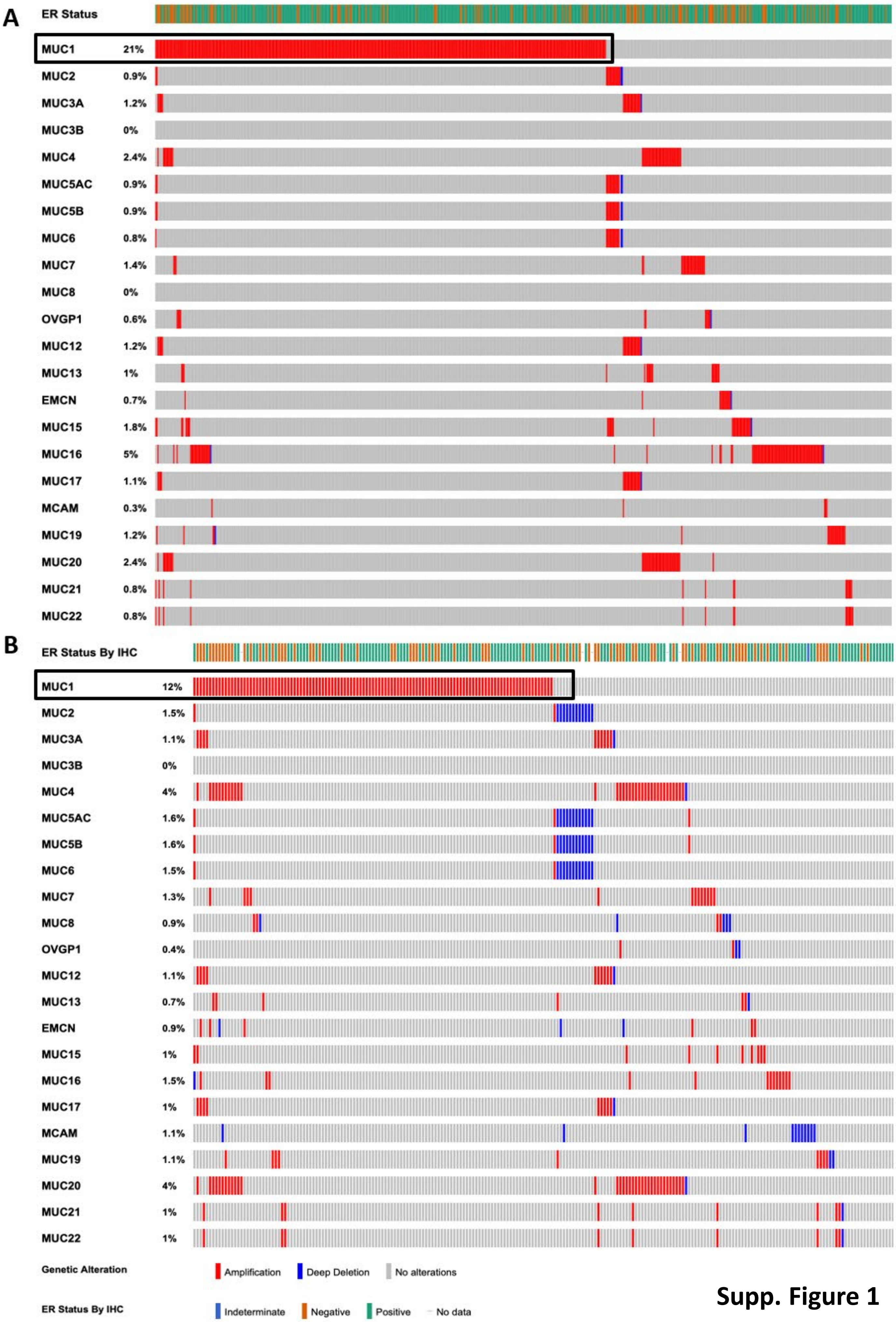


**Supp. Figure 1**

**Supp. Figure 1:** Copy number alterations in mucin family genes in breast cancer tumor samples. OncoPrint visual of genomic alterations on query of mucin family members in METABRIC, n=2509 patients (A) and Firehose Legacy, n=1108 patients (B) using cBioPortal.

### Supp. Figure 2

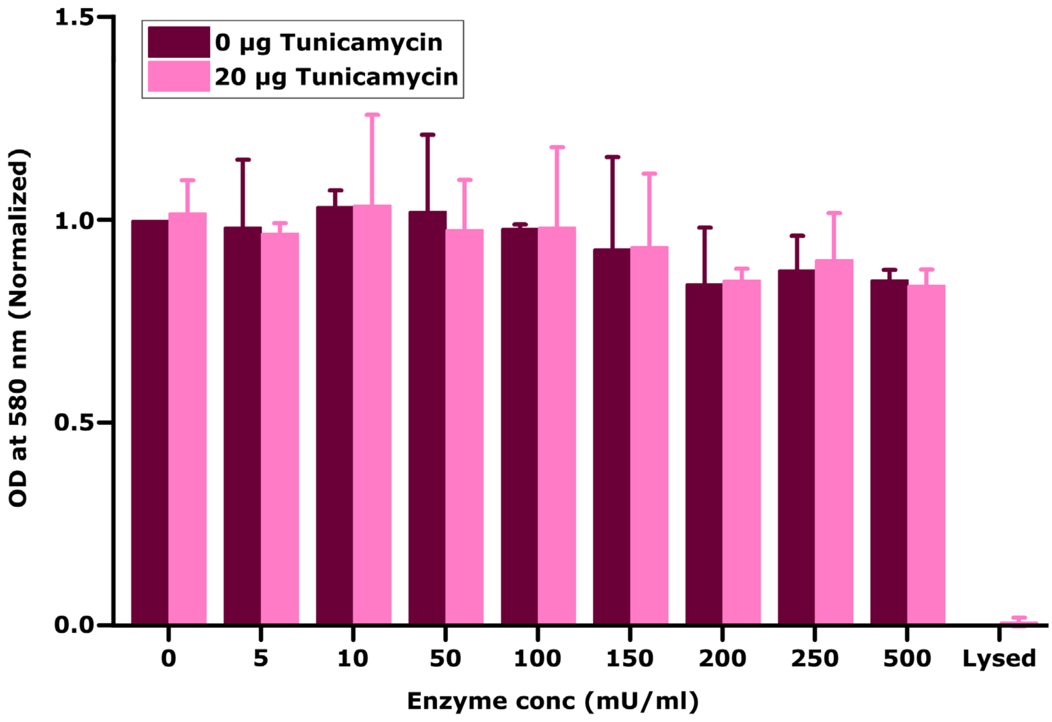


**Supp. Figure 2:** A dose-dependent response with varying concentrations of Neuraminidase and Tunicamycin showing treatment does not significantly affect cell viability. Cells were treated for 48 hrs and MTT assay was done to check viability (n=2, data is presented as mean ± SD).

### Supp. Figure 3

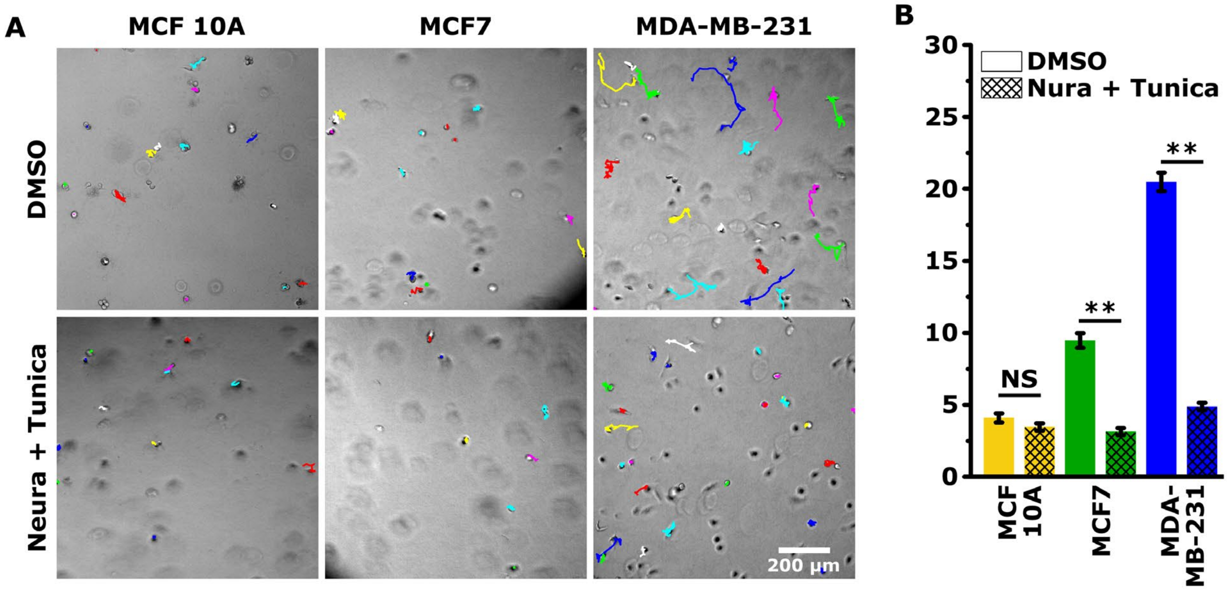


**Supp. Figure 3:** (A) Representative microscopic frames showing cells entrapped in 1mg/ml 3D collagen gel with movement trajectories of MCF 10A, MCF7 and MDA-MB-231 cell in presence of 0.4 U/ml neuraminidase and 20 µg/ml Tunicamycin (NMase) or vehicle (DMSO). (B) Quantification of 3D speed (n=3, per condition >110 cells, ** $p < 0.005$, data is presented as mean ± SEM).

### Supp. Figure 4

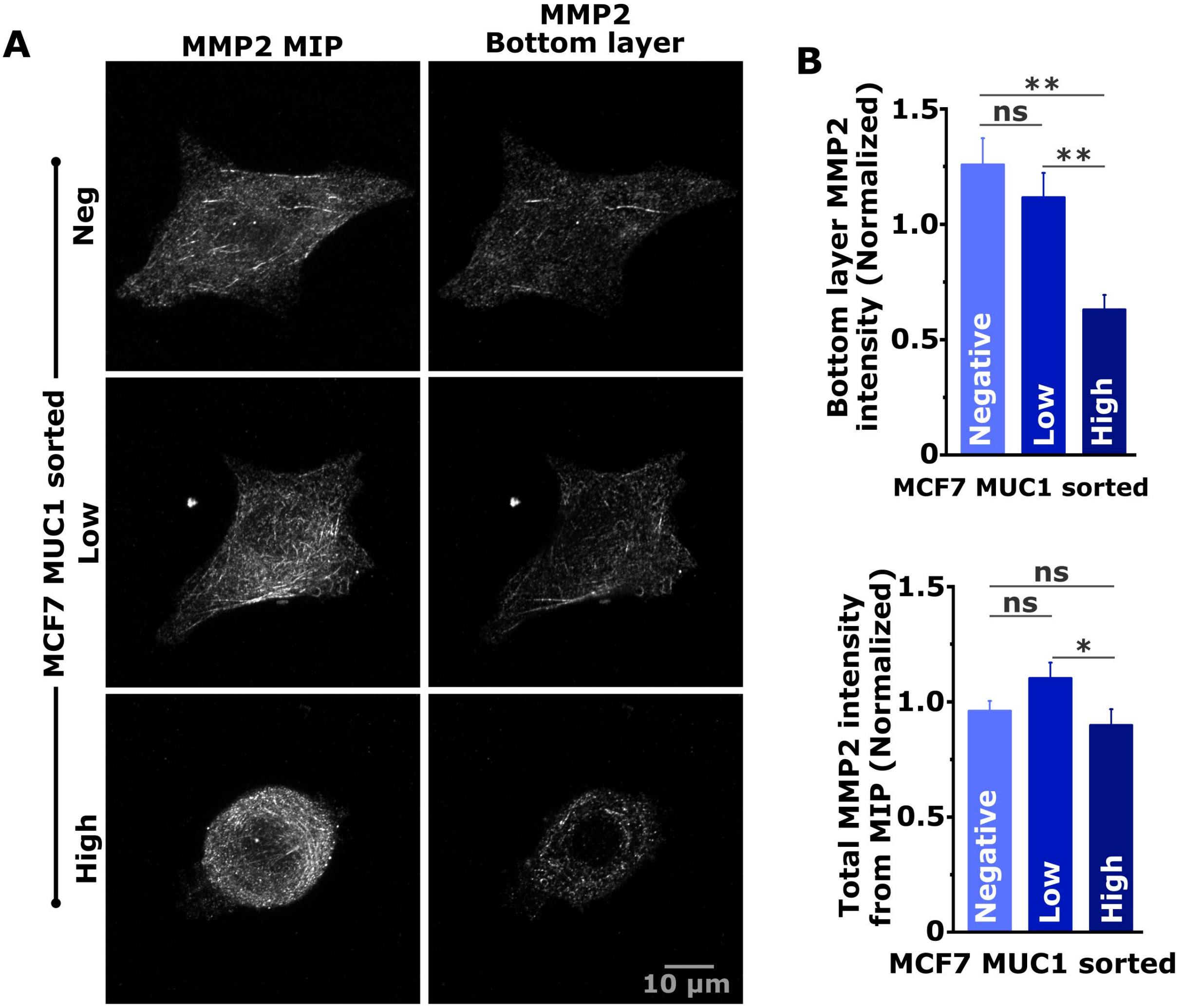


**Supp. Figure 4:** (A) Representative images showing total (maximum intensity projection, MIP) and bottom layer MMP2 levels in MUC1 FACS sorted MCF7 cells cultured on collagen coated substrate. MCF7 cells are sorted in mucin-1 negative, intermediate (low) and high MUC1 expressing cell populations and were grown for 14 hours prior to immunostaining. (B) Quantification of MMP2 expression from obtained image as bottom layer localization (top) and total level from MIP projection(bottom) (n = 3, ** $p < 0.005$, data is presented as mean ± SEM).

### Supp. Figure 5

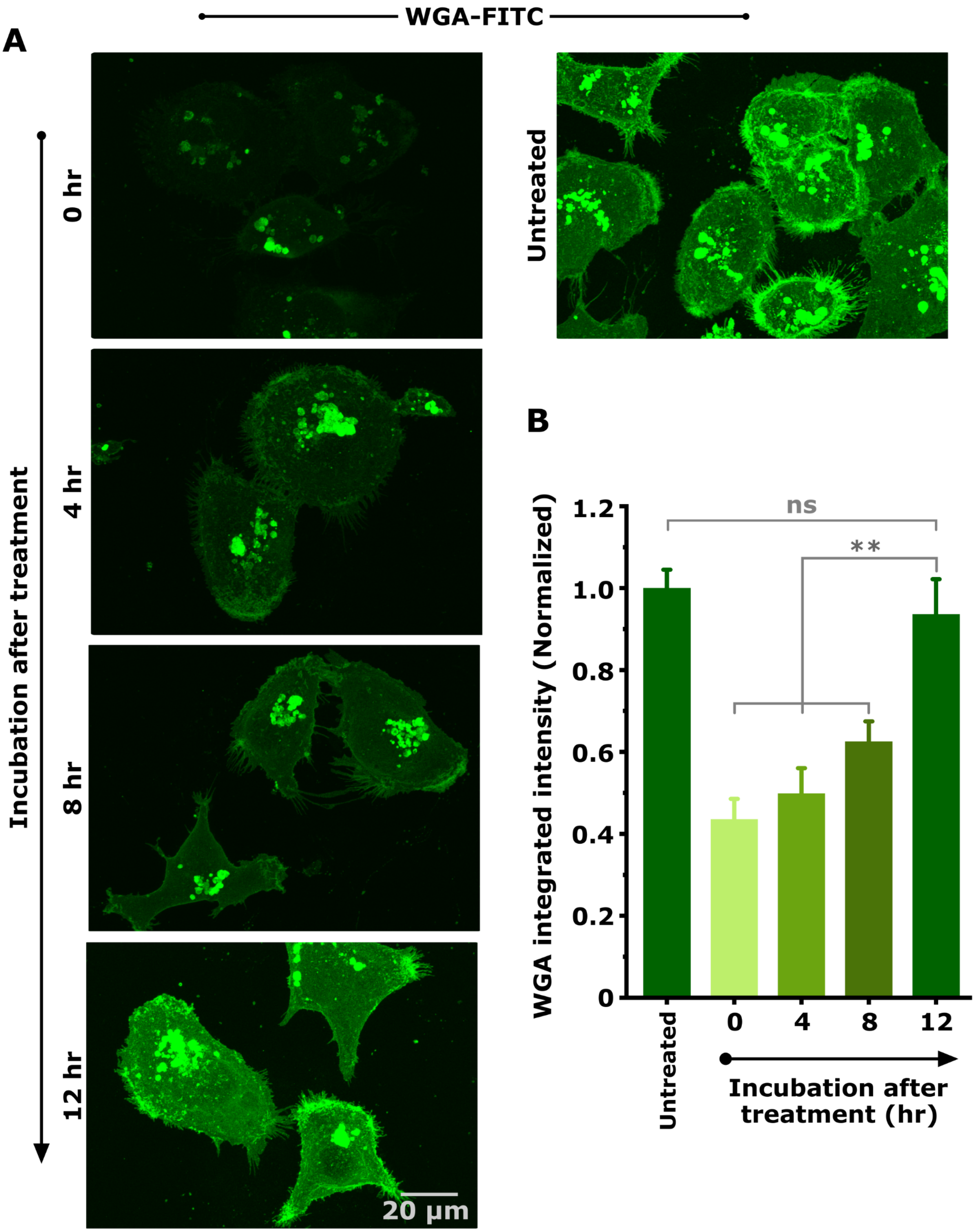


**Supp. Figure 5: Glycan recovery after enzyme treatment.** Cells were cultured for different duration after glycan removal with 0.4 U/ml Neuraminidase treatment to test their surface glycan recovery rate. (A) Representative WGA staining images showing surface glycan levels at different time points after enzyme treatment. (B) Quantification surface glycocalyx recovery rate based on WGA intensity from obtained images (n = 2, ** $p < 0.005$, data is presented as mean ± SEM).

### Supp. Figure 6

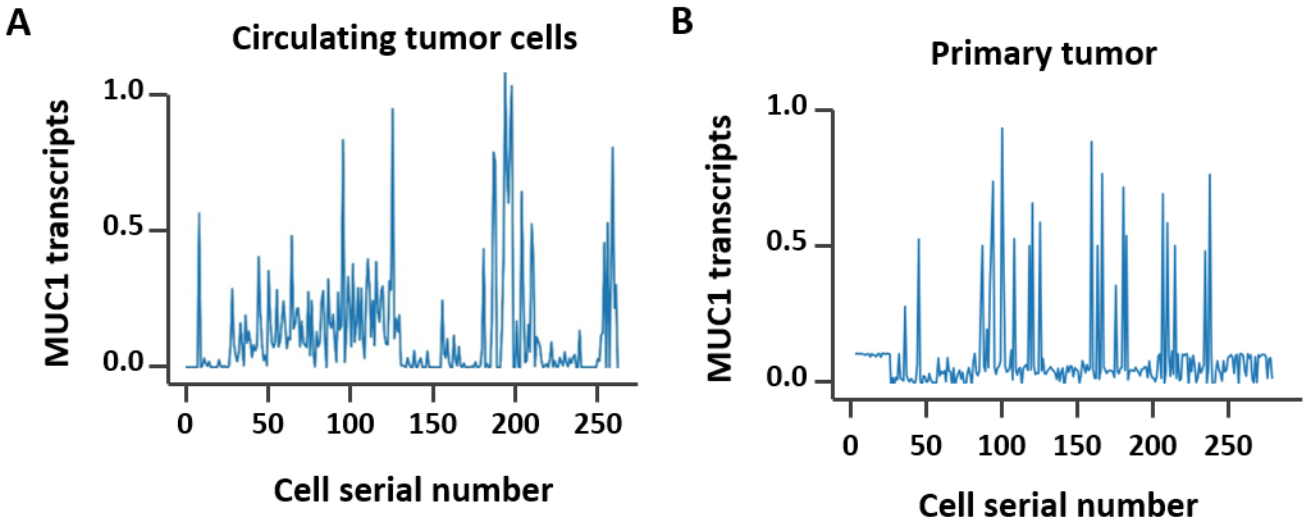


**Supp. Figure 6:** The line plots show the expression profile of the MUC1 gene in patient CTCs (A) and primary tumor cells (B).

**Supp. Movie 1:** FACS sorted MCF7 cells seeded on a selectin coated microfluidics channel were subjected to different flow induced shear stress and were imaged using a phase contrast live cell imaging setup.

**Supp. Movie 2:** WGA staining shows cell leave cell leaves glycan patch after detachment. WGA stained MCF7 cells seeded on a selectin coated microfluidics channel subjected flow induced shear was imaged using fluorescent live cell imaging setup using 60x oil immersion objective.
